# Central oxytocin signaling inhibits food reward-motivated behaviors and VTA dopamine responses to food-predictive cues in male rats

**DOI:** 10.1101/2020.06.24.169540

**Authors:** Clarissa M. Liu, Ted M. Hsu, Andrea N. Suarez, Keshav S. Subramanian, Ryan A. Fatemi, Alyssa M. Cortella, Emily E. Noble, Mitchell F. Roitman, Scott E. Kanoski

**Affiliations:** Neuroscience Graduate Program, University of Southern California, Los Angeles, CA, United States; Department of Biological Sciences, Human and Evolutionary Biology Section, University of Southern California, 3616 Trousdale Parkway, AHF 252, Los Angeles, CA 90089, United States; Department of Psychology, University of Illinois at Chicago, 1007 W. Harrison St., Chicago, IL 60607-7137, United States; Department of Foods and Nutrition, University of Georgia, 129 Barrow Hall, Athens GA 30602, United States

**Keywords:** obesity, impulsivity, incentive, motivation, reward, memory, photometry

## Abstract

Oxytocin potently reduces food intake and is a potential target system for obesity treatment. A better understanding of the behavioral and neurobiological mechanisms mediating oxytocin’s anorexigenic effects may guide more effective obesity pharmacotherapy development. The present study examined the effects of central (lateral intracerebroventricular [ICV]) administration of oxytocin in rats on motivated responding for palatable food. Various conditioning procedures were employed to measure distinct appetitive behavioral domains, including food seeking in the absence of consumption (conditioned place preference expression), impulsive responding for food (differential reinforcement of low rates of responding), effort-based appetitive decision making (high-effort palatable vs. low-effort bland food), and postingestive reward value encoding (incentive learning). Results reveal that ICV oxytocin potently reduces food-seeking behavior, impulsivity, and effort-based palatable food choice, yet does not influence encoding of postingestive reward value in the incentive learning task. To investigate a potential neurobiological mechanism mediating these behavioral outcomes, we utilized in vivo fiber photometry in ventral tegmental area (VTA) dopamine neurons to examine oxytocin’s effect on phasic dopamine neuron responses to sucrose-predictive Pavlovian cues. Results reveal that ICV oxytocin significantly reduced food cue-evoked dopamine neuron activity. Collectively, these data reveal that central oxytocin signaling inhibits various obesity-relevant conditioned appetitive behaviors, potentially via reductions in food cue-driven phasic dopamine neural responses in the VTA.

**Highlights:**

- Central oxytocin inhibits motivated responding for palatable food reinforcement
- Central oxytocin does not play a role in encoding postingestive reward value
- Central oxytocin blunts VTA dopamine neuron activity in response to food cues

## Introduction

There is recent interest in targeting the oxytocin system for obesity pharmacotherapy development based on preclinical findings revealing that oxytocin administration reduces caloric intake, increases fat oxidation, and improves insulin sensitivity (Blevins and Ho, 2013; Lawson, 2017; Olszewski et al., 2017; Sabatier et al., 2013). Oxytocin, synthesized centrally in the paraventricular hypothalamic nucleus (PVH) and supraoptic nucleus (SON), reduces food intake through signaling in classic brain feeding centers, including the arcuate nucleus of the hypothalamus (Caquineau et al., 2006; Fenselau et al., 2017; Maejima et al., 2009; Yosten and Samson, 2010), the ventromedial nucleus of the hypothalamus (Noble et al., 2014), and the hindbrain nucleus tractus solitarius, where oxytocin interacts with other feeding-relevant peptides (i.e. neuropeptide Y, alpha-melanocyte-stimulating hormone, glucagon-like peptide-1 and cholecystokinin) (Affleck et al., 2012; Atasoy et al., 2012; Blevins et al., 2003; Katsurada et al., 2014; Larsen et al., 1997; Motojima et al., 2016; Ong et al., 2015; Ong et al., 2017; Rinaman and Rothe, 2002).

These hypothalamic and hindbrain substrates associated with oxytocin’s feeding effects are thought to predominantly regulate food intake that is driven by energetic need (Liu and Kanoski, 2018). However, feeding is a complex behavior that requires integration of interoceptive and external environmental cues to coordinate food-directed actions, and therefore must also engage “higher-order” substrates that regulate various cognitive processes. Specifically, learned food-predictive cues provide information about properties of impending food availability, signal affective or incentive properties, and elicit skeletal and autonomic responses to guide complex food-directed behavior (Johnson, 2013). While learning about the food environment is necessary for survival, cues associated with palatable food can also stimulate overeating beyond energetic need and contribute to obesity (Ferrario et al., 2016; Hsu et al., 2018a; Johnson, 2013; Kanoski et al., 2013; Liu and Kanoski, 2018; Petrovich and Gallagher, 2007; Rossi and Stuber, 2018). Whether central oxytocin signaling influences higher-order cognitive aspects of feeding that are relevant to excessive caloric intake and obesity is poorly understood. The present study therefore investigated whether central oxytocin signaling influences a multitude of conditioned food-motivated and food cue-associated behavioral tasks that probe [1] palatable food seeking behavior in the absence of consumption, [2] impulsive operant responding for food reinforcement, [3] effort-based decision making (choice between high-effort palatable vs. low-effort bland food), and [4] postingestive reward-value encoding (“incentive learning”).

Conditioned environmental cues that predict the availability of palatable food activate mesolimbic dopaminergic brain reward pathways to enhance food-directed action and consumption (Berridge, 2007; Hsu et al., 2018a). Oxytocin receptors are expressed in dopamine neurons in the ventral tegmental area (VTA) and these oxytocin receptor-expressing VTA neurons project to the nucleus accumbens, prefrontal cortex, and extended amygdala (Peris et al., 2017). Furthermore, oxytocin modulates dopaminergic tone in the VTA and can decrease excitatory synaptic transmission through endocannabinoid-dependent mechanisms (Xiao et al., 2018; Xiao et al., 2017). However, whether central oxytocin signaling affects food cue-associated dopamine signaling in awake behaving animals is unknown. Here we explore the effects of central oxytocin signaling on phasic VTA dopamine neuron responses to sucrose-predictive Pavlovian cues as a potential mechanism contributing to oxytocin’s anorexigenic effects.

## Material and Methods

### Subjects

For experiments 1-5, male Sprague-Dawley rats (Envigo, Indianapolis, IN; weighing >250 g) were individually housed in a temperature controlled (22-23°C) vivarium with ad libitum access to water and chow (LabDiet 5001, Lab Diet, St. Louis, MO; except where noted). Given that some experiments involved chronic food restriction, rats were individually housed for precise control of food rationing. Rats were maintained on a 12hr:12hr reverse light/dark cycle (lights off at 1000h). All procedures were approved by the Institute of Animal Care and Use Committee at the University of Southern California.

For experiment 6, male Long Evans rats expressing Cre recombinase under the control of the tyrosine hydroxylase promoter [TH:Cre+, Rat Research Resource Center, RRRC#:659] were individually housed in a temperature controlled (22-23°C) vivarium, and were moderately food restricted with 15 g of chow per day throughout the duration of their training and experiments. This modest amount of food restriction permitted gradual weight gain throughout training and testing. Rats were maintained on a 12hr:12hr light/dark cycle (lights on at 0700h). Procedures were approved by the Institutional Animal Care and Use Committee at the University of Illinois at Chicago.

### Surgery

For all surgical procedures, rats were anesthetized and sedated via intramuscular injections of ketamine (90 mg/kg), xylazine (2.8 mg/kg), and acepromazine (0.72 mg/kg). Rats were also given analgesic (subcutaneous injection of ketaprofen [5mg/kg] or meloxicam [0.1 mL of 5 mg/mL meloxicam]) after surgery and once daily for 3 subsequent days thereafter). All rats recovered for at least one-week post-surgery prior to experiments.

For central pharmacological oxytocin delivery, rats were surgically implanted with indwelling guide cannula (26-gauge, Plastics One, Roanoke, VA) targeting the lateral ventricle using the following stereotaxic coordinates, which are relative to the location of bregma and dorsal/ventral (DV) coordinates relative to the skull surface at the cannula implantation site: −0.9 mm anterior/posterior (AP), +1.8 mm medial/lateral (ML), and −2.6 mm DV. Cannula were affixed to the skull as previously described using jeweler’s screws and dental cement (Liu et al., 2020). Following one week of recovery, cannula placement was evaluated via measurement of cytoglucopenia-induced sympathoadrenal-mediated glycemic effect that occurs from 210 µg (in 2 µl) of 5-thio-D-glucose (5TG) (Slusser and Ritter, 1980). Food was removed 1 hr prior to the placement test. Rats were injected with 5TG via injectors that extended 2 mm beyond the end of the guide cannula, and blood glucose was measured at 30, 50, 90, and 120 min post-injection. Placement was deemed correct if blood glucose doubled within the measurement period. For rats who did not pass, injector length was adjusted until correct placement was achieved.

For fiber photometry, a Cre-dependent virus containing the construct for a genetically encoded Ca^2+^ indicator (AAV1.Syn.Flex.GCaMP6f.WPRE.SV40) was unilaterally administered to the VTA of TH:Cre+ rats (VTA; 1 µl of 0.5e13 GC/mL: AP -5.4, ML -0.7, DV -815, mm) using a rate of 0.1 µl/min and a 5 min post infusion period to allow for diffusion before the injector was removed. The combination of Cre-dependent construct and transgenic rat leads to strong and selective construct expression in dopamine neurons (Konanur et al. 2020; Decot et al. 2017). Next, an optic fiber (flat 400um core, 0.48 NA, Doric Lenses Inc.) was implanted directly above the injection site (AP -5.4, ML -0.7, DV -8.00). Lastly, an ICV cannula was implanted as described above. Rats were given 2-week recovery prior to behavioral training procedures. Rats with missed fiber optic placement were excluded from the experiment (n=3 removed).

### Behavior

#### Conditioned Place Preference (CPP)

CPP training and testing procedures (Fig 1A) were conducted as described previously (Hsu et al., 2015; Kanoski et al., 2014b; Kanoski et al., 2011b). Briefly, the CPP apparatus consisted of two conjoined plexiglass compartments with a guillotine door separating the two sides (Med Associates, Fairfax, VT, USA). The two sides (contexts) were distinguished by wall color and floor texture. Rats (n=18, n=8/aCSF, n=9/oxytocin) were given one 15 min habituation session with the guillotine door open and video recording to measure time spent in each context. For each rat, the least preferred context during habituation was designated as the food-paired context for subsequent training.

**Figure 1:**
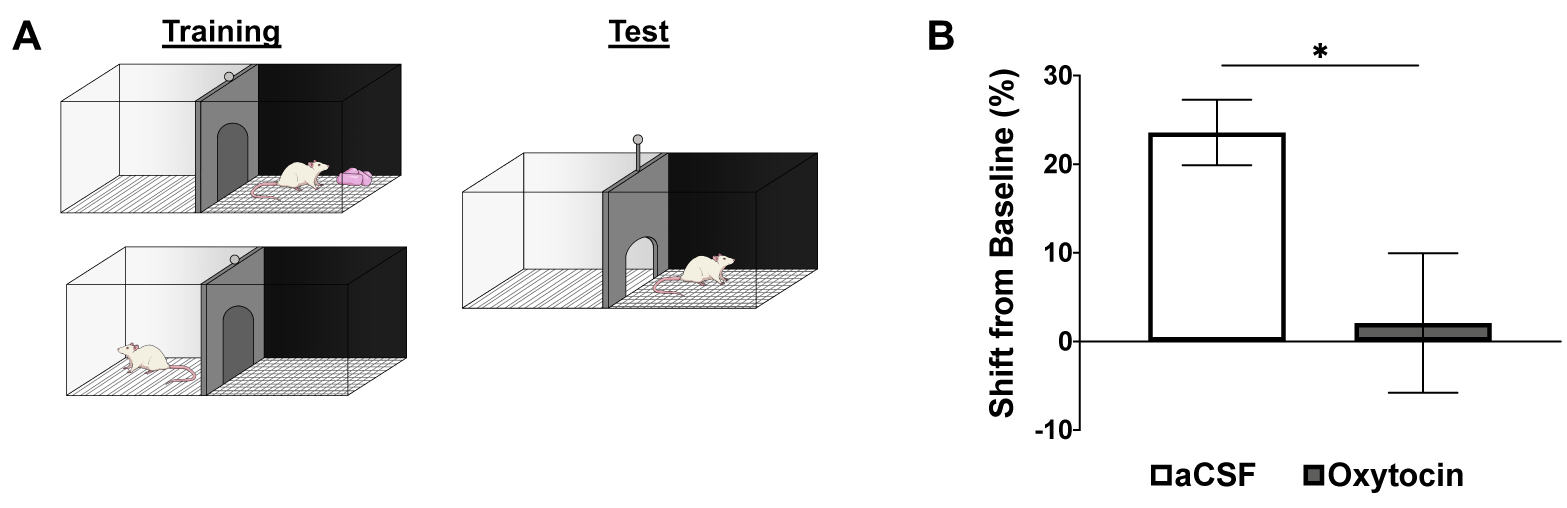
ICV oxytocin suppresses food seeking behavior in the conditioned place preference (CPP) test. (A) Schematic of CPP training and test procedures. (B) Effect of ICV oxytocin on time spent on the food-paired side during the test (represented as a percent shift from baseline). Data are mean ± SEM; *p<0.05.

Training occurred in the early phase of the dark cycle and home-cage chow was pulled 2.5 hr prior to each training session. CPP training consisted of 16 daily, 20-min sessions: eight sessions isolated in the food-paired context and eight sessions isolated in the non-food-paired context. Context training order was randomized. During the food-paired sessions, 5 g of 45% kcal high fat/sucrose diet (D12451, Research Diets, New Brunswick, NJ, USA) was placed on the chamber floor, and no food was presented during non-food-paired sessions. All rats consumed the entire 5 g of food during each food-paired session.

CPP testing occurred 2 days after the last training session using a between-subjects design, and groups were matched for baseline context preference. Immediately before the testing session, rats received either lateral ICV 1 µg/1µl oxytocin (Bachem, Torrance, CA, USA) or artificial cerebrospinal fluid (aCSF) vehicle injections. During testing, the guillotine door remained open and rats were allowed to freely explore both contexts for 15 min. No food was present during testing. Time spent in each context during the test was later calculated from video recordings by an experimenter blind to the group and context-food assignments. The dependent variable was the percentage shift in preference for the food-associated context during testing compared with the baseline session.

#### Differential reinforcement of low rates of responding (DRL)

DRL training and testing procedures (Fig 2A) were conducted as described previously (Hsu et al., 2018b; Noble et al., 2019). Rats (n=12, within-subjects design) were first habituated in their home cage to consume twenty 45 mg sucrose pellets (F0023, Bio-Serv, Flemington, NJ, USA)). Throughout DRL training (5 days/week), home cage chow was removed 1 h prior to training, which began at the onset of the dark cycle, and was returned to the rats following training. During training, rats were placed in an operant chamber (Med Associates, Fairfax, VT, USA) containing an active lever (reinforced with one 45 mg sucrose pellet) and an inactive lever (non-reinforced). Training sessions were 45 min long, during which both levers were extended at the start of the session and retracted at the end of the session. Neither lever retracted at all during the session, and no visual or audio cues were used during training or testing. For the first five days of training, rats were on a DRL0 schedule, where each active lever press was reinforced with a 0 sec time delay. They were then switched to a DRL5 schedule for five days, where rats had to withhold presses on the active lever for at least a 5 sec interval for each active lever press to be reinforced. Active lever presses that occurred before the 5 sec had elapsed were not reinforced and the timer was restarted. They were then switched to 5 days of DRL10 (10 sec withholding period) and 10 days of DRL20 (20 sec withholding period). Efficiency in DRL was calculated as the number of pellets earned divided by the number of active lever presses.

**Figure 2:**
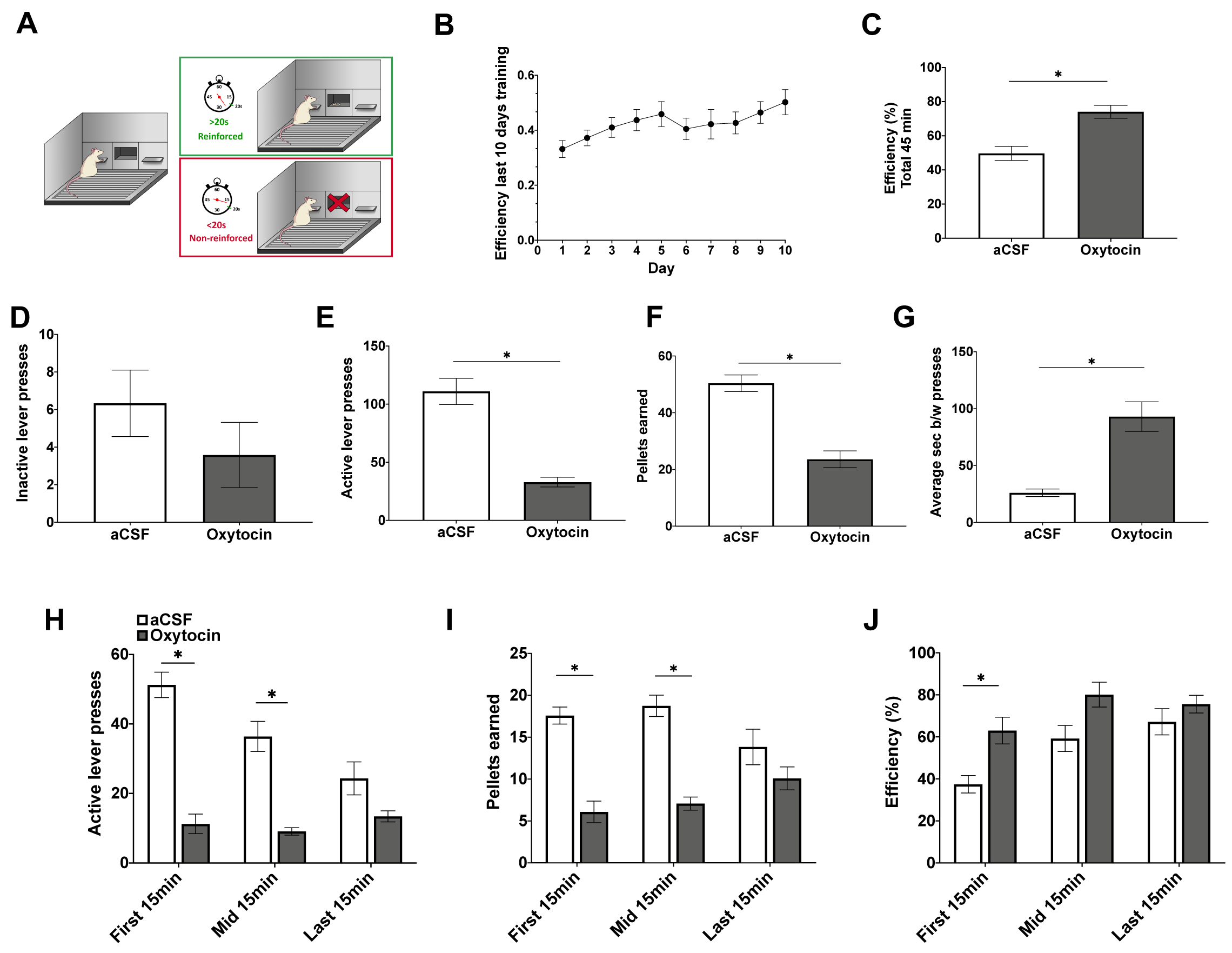
ICV oxytocin reduces impulsive responding in the differential reinforcement of low rates of responding (DRL) task. (A) Schematic of DRL training and testing procedures. (B) Efficiency (pellets earned divided by number of active lever presses) over the last 10 days of DRL20 training. (C) Efficiency for entire test session (45 min) following ICV administration of oxytocin. (D-F) Total inactive lever presses, active lever presses, and sucrose pellets earned for entire test session following ICV administration of oxytocin. (G) Average seconds between active lever presses following ICV administration of oxytocin for entire test session. (H-J) Active lever presses, pellets earned, and efficiency broken into 15 min time blocks following ICV administration of oxytocin. Data are mean ± SEM; *p<0.05.

The DRL test was conducted using a within-subjects design and no training sessions occurred on days between tests. Oxytocin (Bachem, Torrance, CA, USA) or vehicle treatment order was counterbalanced based on efficiency during training, and treatments were separated by 72 h. On test days, home cage chow was removed 1 h prior and testing began at dark onset. ICV injections of 1 µg/µl oxytocin or vehicle were administered immediately prior to testing on a DRL20 schedule.

#### Progressive Ratio (PROG) / Chow Feeding Choice Task

Training and testing in the PROG/Chow choice task (Fig 3A) was adapted from Salamone and colleagues (Randall et al., 2012; SanMiguel et al., 2018; Yohn et al., 2016). Following surgical recovery, rats (n=11, within-subjects design) were brought down to and maintained at 85% of their free-feeding body weight. Rats were habituated to twenty 45 mg sucrose pellets (F0023, Bio-Serv, Flemington, NJ, USA) in the home cage prior to training. Behavioral sessions (30 min, 5 days/week) were conducted in operant conditioning chambers (Med Associates, Fairfax, VT, USA) and began at the onset of the dark cycle. Rats were initially trained to lever press on a FR1 schedule (one active lever press for one sucrose pellet) for 5 days, and then shifted to a PROG schedule for 10 days. During the PROG session, the ratio started at FR1 and sequentially increased by one every time 15 reinforcers were obtained (FR1 × 15, FR2 × 15, FR3 × 15….). The rats were then introduced to free chow (Laboratory Rodent Diet 5001, St. Louis, MO, USA) concurrently available in addition to the PROG sessions for 5 days. Active lever presses, inactive lever presses, pellets earned, highest PROG ratio achieved, and chow intake (including spill) were recorded throughout training.

**Figure 3:**
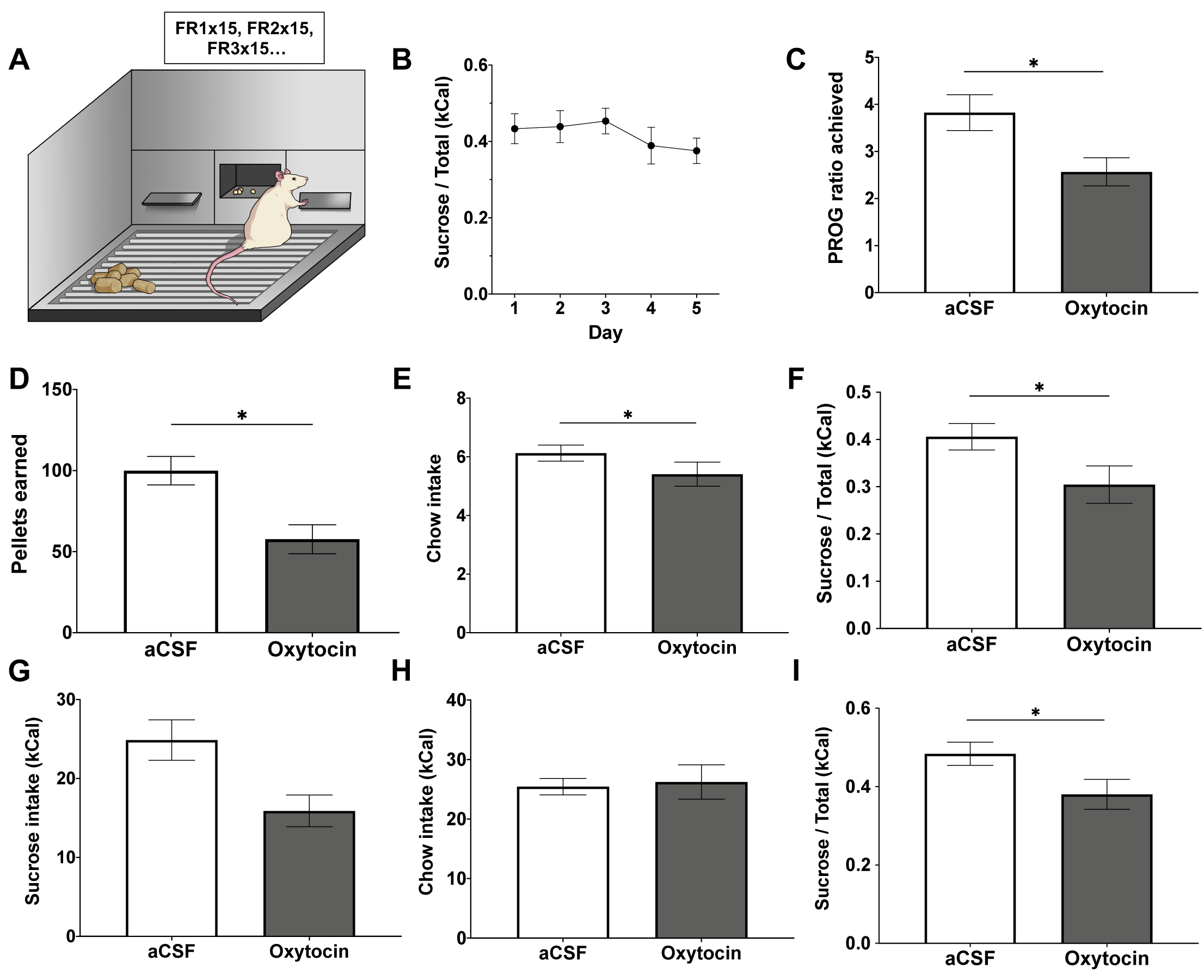
ICV oxytocin reduces effort-based consumption of sucrose over freely available chow in a PROG/Chow choice task. (A) Schematic of the PROG/Chow choice task. (B) Ratio of sucrose intake over total intake (kCal) over the course of five days training in the choice task. (C-F) Highest PROG ratio achieved, PROG sucrose pellets earned, free chow intake, and ratio of PROG sucrose intake over total intake in the PROG/Chow choice task following ICV oxytocin administration. (G-I) Free sucrose pellets consumed, free chow intake, and ratio of free sucrose intake over total intake in free sucrose/free chow choice task following ICV oxytocin administration. Data are mean ± SEM; *p<0.05.

Testing in the PROG/Chow feeding choice task was conducted using a within-subjects design, and treatments were separated by 72 h. Drug treatment order was counterbalanced and matched by performance during the last day of training (ratio of sucrose to chow intake, kCal). The rats received lateral ICV 1 µg/µl oxytocin (Bachem, Torrance, CA, USA) or vehicle immediately prior to 30 min test session, where PROG performance and chow intake were measured (just as in training sessions).

In addition to the PROG/Chow feeding choice task, the effect of ICV oxytocin on preference between consumption of free chow and free sucrose concurrently available in the apparatus was also examined (n=10, within-subjects design). Rats were maintained at 85% of their free-feeding body weight and tested at the onset of the dark cycle. Treatments were separated by 48 h and drug treatment order was counterbalanced. Using the same apparatus (but without extension of the levers) and drug injection parameters as the PROG/Chow feeding choice task, intake of freely available chow and freely available sucrose was measured in a 30 min test session and ratio of intake was calculated.

#### Incentive Learning

Rats were trained in an instrumental incentive learning task, modified from (Wassum et al., 2011a; Wassum et al., 2011b; Wassum et al., 2009). Rats (n=21, n=6/dep aCSF, n=8/dep OT, n=8/sated aCSF) were habituated to twenty 45 mg sucrose pellets (F0023, Bio-Serv, Flemington, NJ, USA) in the home cage prior to training. Training sessions occurred daily at the onset of the dark cycle, and food was pulled 1 hr prior to the start of the session and returned 1 hr after completion of the session. Each session started with insertion of the levers where appropriate and ended with retraction of the levers.

For magazine and single-action instrumental training, rats received 3 d of magazine training where they received 20 noncontingent sucrose deliveries in the operant chamber with the levers retracted. Following magazine training, rats received 3 d of single-action instrumental training, where right lever presses were continuously reinforced up to 20 pellets or 30 min elapsed.

Rats were next trained in the reward-delivery chain phase (Fig 4A). Initially, only the left (seeking) lever was retracted. One left lever press was rewarded by presentation of the right (taking) lever, and one right lever press was rewarded with delivery of one 45 mg sucrose pellet. This reward-delivery chain lasted until 20 pellets were earned or 30 min elapsed. The seeking lever was continuously rewarded with the taking lever for four sessions, then the reinforcement schedule was increased to random ratio (RR)-2 for four sessions, and then to RR-4 until stable lever pressing was obtained (approximately three to five sessions). The taking lever was always continuously rewarded (FR-1) and retracted after sucrose delivery. Rats that did not earn 20 sucrose pellets on a RR-4 schedule were pulled out from the experiment.

**Figure 4:**
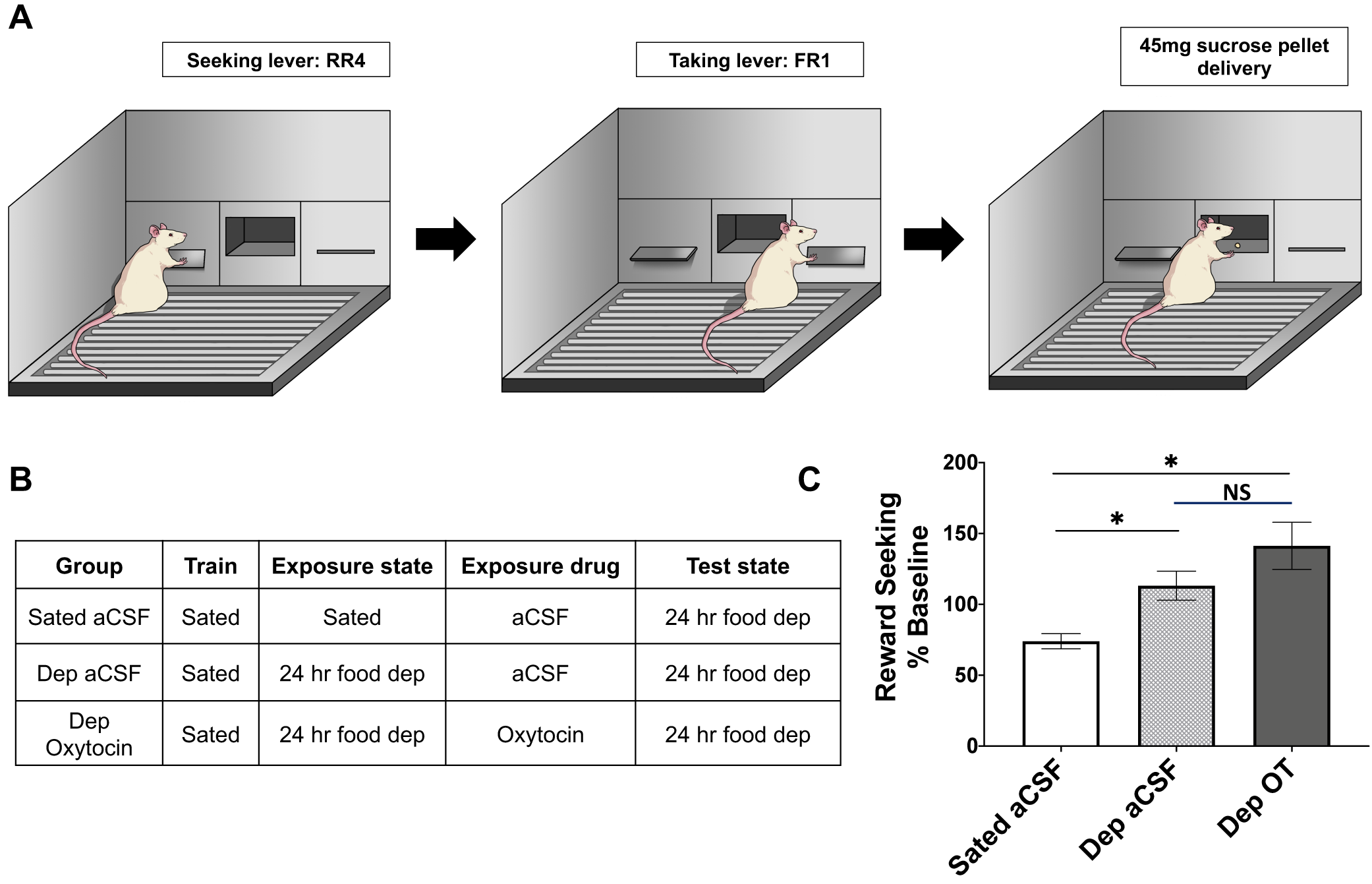
Oxytocin does not play a role in postingestive reward value encoding in an instrumental incentive learning task. (A) Schematic of reward-delivery chain in an instrumental incentive learning task. (B) Table outlining drug treatment and deprivation state for three experimental groups: sated aCSF, 24 hr food deprived (dep) aCSF, 24 hr food deprived (dep) oxytocin. (C) Reward seeking behavior (% baseline) on the reward delivery chain represented by rate of presses on the right (taking) lever during 4 min extinction session. All rats tested under 24 hr food deprivation. Data are mean ± SEM; *p<0.05.

For the US exposure phase, rats were then split into groups of 1 h restriction (sated) + aCSF, 24 h restriction (dep) + aCSF, or 24 h restriction (dep) + oxytocin (Fig 4B). Rats were appropriately food restricted and received 1 µg/µl oxytocin or vehicle ICV injections immediately prior to an exposure session. Exposure session was held in a novel context, where all rats received 30 noncontingent sucrose pellets to consume in 40 min. The exposure session was followed by 2 d of ad libitum re-feeding in the home cage.

For the incentive learning test session, all rats were 24 h food-deprived prior to the test session that occurred in the operant chambers. Rats were tested under extinction in the reward-delivery train on an RR-4 reinforcement schedule for 4 min. Right lever presses, left lever presses, and seconds between right (taking) lever presses were measured.

#### Open field

General activity level and anxiety were measured in open field, which consists of a large gray bin, 60 cm (L) X 56 cm (W), that is placed under diffuse even lighting (30 lux). A center zone is marked in the bin [19 cm (L) X 17.5 cm (W)]. AnyMaze Software (Stoelting Co., Wood Dale, IL) was used to track rat movement. Rats (n=9, within-subjects) were tested under ad lib feeding conditions 1 hr after the onset of the dark cycle. Rats received 1 µg/µl oxytocin or vehicle ICV injections immediately prior to the test, where they were allowed to freely explore the arena for 10 min. The apparatus was cleaned with 10% ethanol after each rat was tested.

#### Cue-predictive reward task for measurement of VTA dopamine neuron activity

After in vivo fiber photometry preparation, food restricted rats (n=4) were trained to associate a cue with brief access to sucrose. Training and experimental sessions were conducted during the light phase in standard operant chambers (ENV-009A-CT, Med Associates, Fairfax, VT, USA). While the procedures described above were conducted during the dark cycle, photometry protocols were established and optimized in the light cycle (Konanur et al., 2020) and thus rats underwent daily training for 8 consecutive days during the light cycle. Each training session was comprised of 40 trials where a trial consisted of a 1 sec audio cue (4.5 kHz tone or white noise) followed by either 20 sec availability of a retractable sipper containing a 0.3 M sucrose solution (CS+ trials; n=20) or a dry sipper (CS-trials; n=20). Licks were timestamped using a contact lickometer and controller (ENV-252 M; ENV-250, Med Associates Inc.). The audio cue paired with each sipper was counterbalanced across rats. Trials were separated by a randomly selected, variable inter-trial interval (32-48 sec). Following eight training days, test sessions were conducted identical to training sessions but lateral ventricle injections of vehicle, 0.3 µg/ µl oxytocin, or 1 µg/µl oxytocin (within-subjects) were made immediately prior.

### Fiber Photometry and Signal Normalization

For in vivo fiber photometry, LEDs delivered 465 nm (Ca^2+^ dependent) and 405 nm (Ca^2+^ independent) excitation. Intensity of the 465 nm and 405 nm light were sinusoidally modulated at 211 Hz and 531 Hz, respectively, for all recording sessions. Intensity was then coupled to a filter cube (FMC4 contains excitation filters at 405 nm, 460–490 nm and emission filters at 500–550 nm, Doric Lenses), and converged into an optical fiber patch cord mated to the fiber optic implant. GCaMP6f fluorescence was collected and focused onto a photoreceiver (Visible Femtowatt Photo-receiver Model 2151, Newport). A lock-in amplifier and data acquisition system (RZ5P; Tucker Davis Technologies) was used to demodulate 465 nm and 405 nm excited fluorescence. Behavioral events (e.g. cue, licks) were timestamped and sent as digital inputs to the same data acquisition system and recorded in software (Synapse Suite, Tucker Davis Technologies). The signal was then processed with Fourier transformed subtraction to calculate fluorescence due to fluctuations in Ca^2+^ and bleaching and movement artifacts (ΔF/F). The subtracted signal was smoothed using a custom fifth order bandpass butterworth filter (cutoff frequencies: 0.05 Hz, 2.25 Hz).

To compare behavior-related responses across recording sessions, the smoothed Fourier subtracted Ca^2+^ specific signal of each session was normalized by the mean transient Ca^2+^ amplitude of that session (Konanur et al., 2020). A transient was defined as a point 3 standard deviations above the previous point. The normalized signal was then aligned to a 15 sec cue window around cue onset. All data processing was performed using custom MATLAB scripts (available upon request).

### Statistical Analyses

All experiments met assumptions of normality (Shaprio Wilk), and Experiments 2-6 met assumptions for equality of variance (Levene’s tests). Experiment 1 did not pass assumption of equality variance test, and was thus analyzed using between-subjects Welch’s t-test. Experiments 2, 3, and 5 were analyzed using paired t-tests. 15-min time blocks of active presses, pellets earned, and efficiency in experiment 2 (DRL) were analyzed using two-way analysis of variance (ANOVA) with time and drug as within-subjects variables, with Bonferroni post-hoc tests for multiple comparisons. Experiment 4 was analyzed using a one-way ANOVA and Bonferroni post-hoc tests for multiple comparisons.

All photometry data were analyzed using parametric repeated measures one or two-way ANOVAs. In all photometry experiments, signal windows were averaged across trials for individual rats, then between rats. For experiment 5, acquisition of CS+/CS-discrimination was measured by 1^st^ lick latency and % of trials containing at least one lick. Analyses here utilized two-way repeated measures ANOVA with cue and day as within-subjects variables. The magnitude of cue-evoked dopamine neuron transients (mean of signal 0 s to 1 s after cue onset) during CS+ compared to CS-were analyzed using student’s t-test. In oxytocin treatments involving fiber photometry, mean signal 1 s after cue onset and behavioral measures in each drug treatment were compared using one-way repeated measures ANOVAs. When drug main effects were found, Bonferroni post-hoc tests were used to compare individual drug treatments.

Effect size estimates were calculated for all significant group differences. Statistical analyses were performed using Graphpad Prism 8 with an α level of significance of p < .05.

## Results

### Experiment 1: Central oxytocin reduces food seeking behavior in the absence of consumption (conditioned place preference expression)

ICV oxytocin administration prior to testing significantly decreased conditioned place preference expression for the palatable food-associated context relative to vehicle (expressed as % shift from baseline preference, Fig 1B) (t_[15]_ = 2.47, p = 0.03; effect size d = −1.151), suggesting that central oxytocin signaling reduces palatable food seeking behavior in the absence of consumption.

### Experiment 2: Central oxytocin decreases food impulsivity (differential reinforcement of low rates of responding task)

ICV administration of oxytocin significantly reduced impulsive behavior in the DRL20 schedule (Fig 2). Oxytocin significantly increased the average number of seconds between active lever presses (t_[11]_ = 6.17, p = 7E-05, fig 2G). Vehicle-treated rats waited an average of 26.05 sec between active lever presses, whereas ICV oxytocin-treated rats waited an average of 92.95 sec between active lever presses. Furthermore, ICV oxytocin significantly decreased the number of sucrose pellets earned (t_[11]_ = 9.17, p = 1.7E-06, fig 2F) and active lever presses (t_[11]_ = 9.04, p = 2E-06, fig 2E) in the DRL test. There were also no significant differences in number of inactive lever presses (fig 2D). Overall efficiency during the test (fig 2C), a measure of impulsive behavior, was calculated by dividing pellets earned by number of active lever presses. ICV oxytocin treatment significantly increased efficiency in the task (t_[11]_ = 5.11, p = 0.0003, fig 2C; effect size d = 1.759), indicating reduced impulsivity. When efficiency is broken down into three 15 min time blocks, there is a significant time (p < 0.0001) and drug (p = 0.0075) main effect, but no significant time by drug interaction. However, post-hoc analyses reveal significant differences in efficiency between oxytocin and vehicle treatments in the first 15 min, but not the mid 15 min or last 15 min (fig 2J).

### Experiment 3: Central oxytocin shifts food choice from high-effort palatable towards low-effort bland food (PROG versus chow choice task)

Motivated behaviors are often characterized by increased work to obtain reinforcement, so we tested effort-based decision making using the PROG versus choice task. A 2-treament within-subjects analysis demonstrates that oxytocin significantly decreased highest PROG ratio achieved (t_[10]_ = 5.63,p = 0.0002, fig 3C), pellets earned (t_[10]_ = 7.76, p < 0.0001, Fig 3D), and free chow consumption (t_[10]_ = 2.51, p = 0.03, fig 3E). Moreover, oxytocin significantly decreased the ratio of sucrose over total intake (t_[10]_ = 4.29, p = 0.0016; effect size d = −0.889, Fig 3F), suggesting a preferential decrease in motivation for high-effort sucrose in favor of low-effort chow. In addition to examining effort-based choice, the effect of oxytocin on preference between freely available chow and freely available sucrose in the conditioning chamber was also measured. ICV oxytocin significantly reduced sucrose intake (t_[9]_ = 3.77, p = 0.0045, Fig 3G). Interestingly, ICV oxytocin had no effect on chow intake (Fig 3H). Thus, oxytocin significantly reduced the ratio of sucrose intake relative to chow intake (t_[9]_ = 2.37, p = 0 0.042; effect size d = −1.412, Fig 3I).

### Experiment 4: Central oxytocin does not affect postingestive reward value encoding (incentive learning task)

Incentive learning is the process of modifying the incentive value of a specific food through outcome-nutritive state learning (Balleine, 2001; Balleine and Dickinson, 1998a; Balleine and Dickinson, 1998b). Results reveal that ICV oxytocin did not influence the capacity of food restriction to update/enhance the incentive value of consuming sucrose. Analysis of reward seeking behavior during testing (as a percent of baseline) revealed a significant group effect (F_[2,16]_ = 4.52, p = 0.005; effect size d = 1.24). Posthoc comparisons revealed that the dep aCSF and dep oxytocin groups were significantly different from the sated aCSF group, but not from each other.

### Experiment 5: Central oxytocin administration does not affect general activity levels or anxiety-like behavior in an open field test

ICV oxytocin or vehicle was administered immediately prior to an open field test, where rats are allowed to freely move in a novel chamber devoid of food or food-related cues. Results reveal that ICV oxytocin did not significantly affect the distance traveled (Fig 5A), nor did it affect time spent in the center zone (Fig 5B).

**Figure 5:**
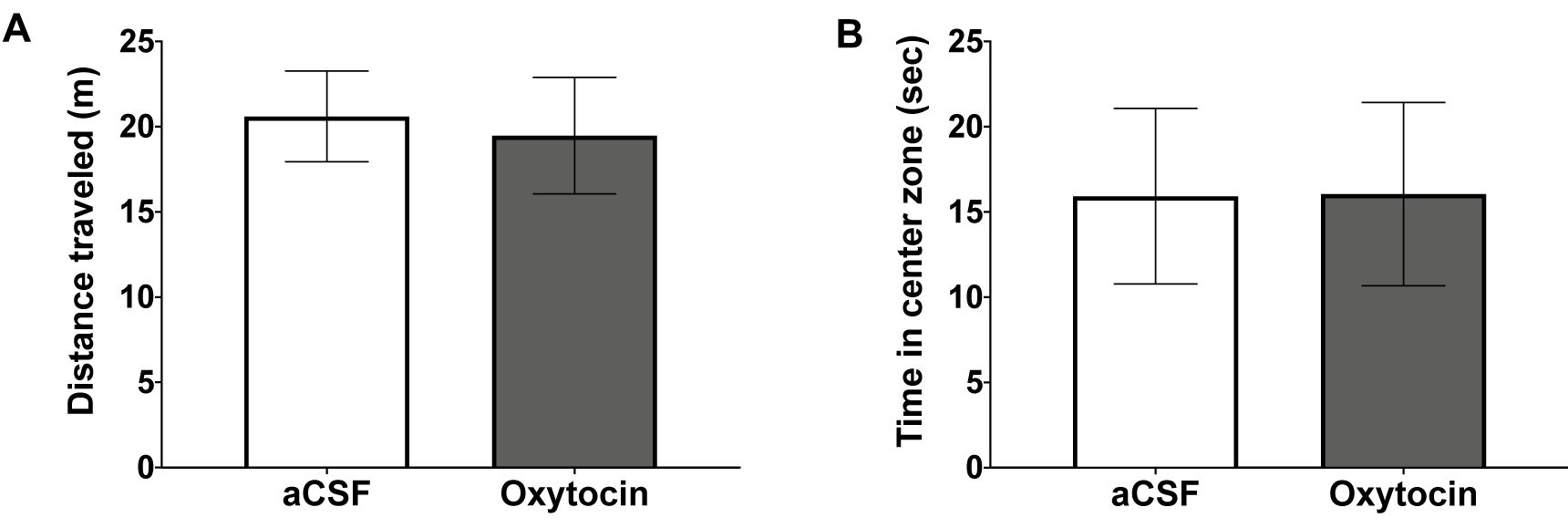
ICV oxytocin does not affect locomotion or anxiety-like behavior in the open field test. (A) Distance traveled in the open field following ICV oxytocin. (B) Total time spent in the center zone in the open field following ICV oxytocin.

### Experiment 6: Central oxytocin administration suppresses VTA phasic dopamine neuron activity in response to cues associated with palatable food

Rats were trained to discriminate between a conditioned stimulus plus (CS+) that predicted the presentation of sucrose reward and a conditioned stimulus minus (CS-) that predicted no reinforcement. As training progressed, rats successfully discriminated between CS+ and CS-cues as measured by significant differences in latency to first lick and percent of trials with a lick (Fig 6A (top): F_[7, 21]_ = 5.30, p = 0.0013 main effect of day; F_[1, 3]_ = 212.77, p = 0.0007 main effect of cue; F_[7, 21]_= 19.95 p < 0.0001 day x cue interaction. Fig 6A (bottom): F_[7,21]_ = 1.405 p = 0.26 no main effect of day; F_[1, 3]_ = 253.50 p = 0.0005 main effect of cue; F_[7,42]_ = 17.06 p<0.0001 day x cue interaction). Fiber photometry (Fig 6B, histology and fiber optic placement in Fig 6C-D) revealed a robust, time-locked dopamine response to the CS+, but not the CS-(Fig 6E; t(3) = 8.49 p = 0.0034). Further, differences between the CS- and CS+ dopamine response 1 sec post-cue had an (effect size d = 1.025). We found no effect of central oxytocin on cue-evoked VTA phasic dopamine activity across the entirety of the behavioral session (veh: 0.22±0.05; 0.3µg OT: 0.18±0.03; 1µg OT: 0.18±0.05; data are mean±SEM. F_[2,6]_ = 2.089, p = 0.21). However, a priori analyses were also conducted across the 1st 10 trials (minutes 0-20) based on the short half-life of central oxytocin (Mens et al., 1983) and based on our impulsivity data revealing maximum drug effect in the 1st 15-min epoch (Fig 2H-J). Results from this analysis revealed that both the 0.3 µg and 1 µg dose significantly reduced cue-evoked VTA phasic dopamine responses relative to vehicle treatment (Fig 6F, 6G; F_[2,6]_ = 8.880 p= 0.016 main effect of drug; Bonferroni post-hoc: p = 0.036 vehicle vs. 0.3 µg oxytocin, p = 0.015 vehicle vs. 1 µg OT). Additionally, we demonstrate that central oxytocin has no impact on 1^st^ lick-evoked VTA phasic dopamine activity (Fig 6H, Ca2+ signal aligned to first lick, F[2,6]=0.17, p=0.85, no main effect of treatment), suggesting that central oxytocin signaling impacts VTA phasic dopamine responses to cues associated with food, but not consummatory behaviors.

**Figure 6:**
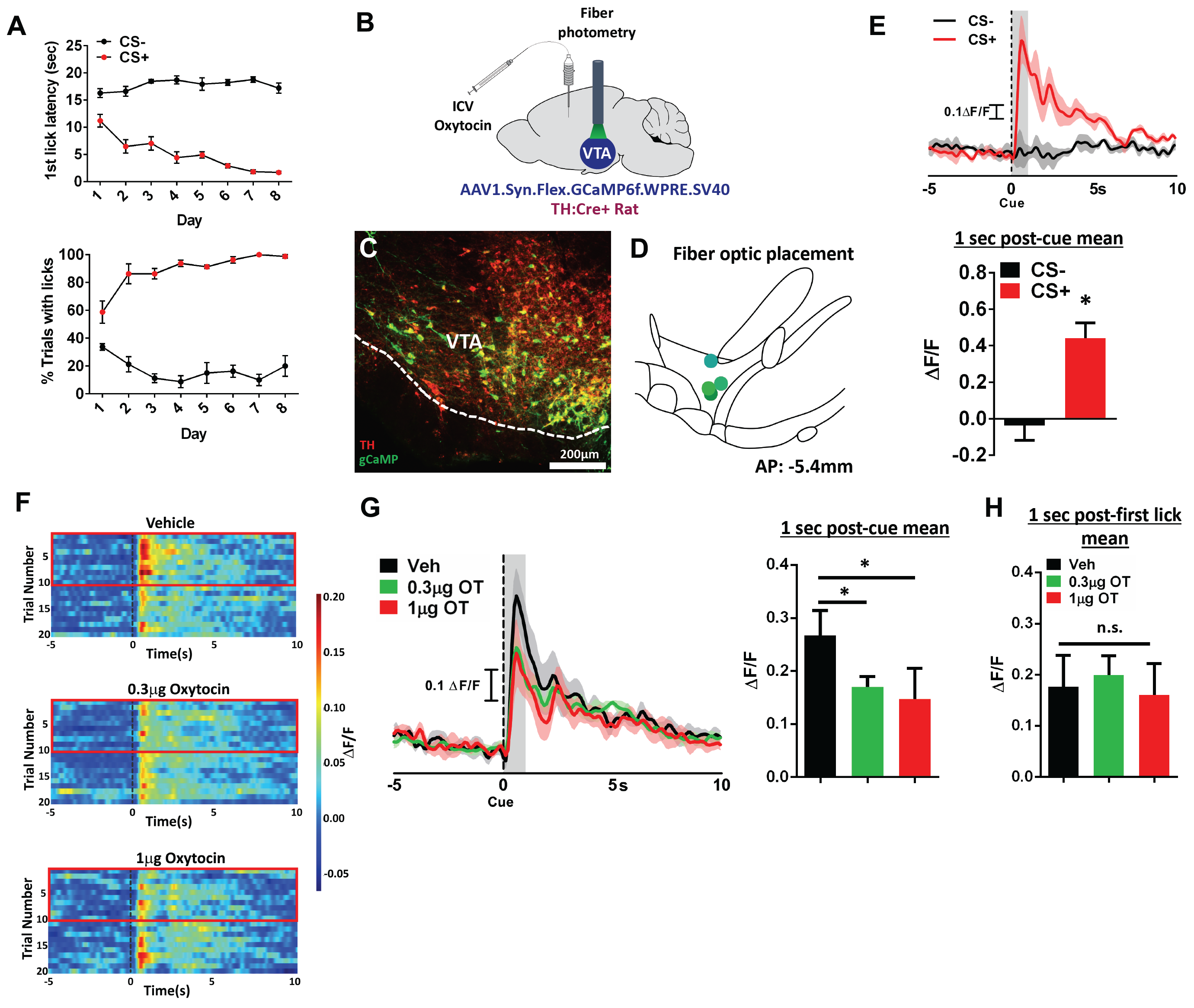
ICV oxytocin reduces food cue-evoked dopamine neuron response in the ventral tegmental area. (A) First lick latency and % trials with licks following the CS+ and the CS-during training, where CS+ precedes sucrose presentation and CS-precedes no food reward. (B) Experimental schematic of fiber photometry and oxytocin pharmacology in TH:Cre+ rats (C) Representative AAV1-FLEX-gCaMP (green) and tyrosine hydroxylase (red) expression in the VTA. (D) VTA fiber optic placement. (E) Representative trace (top) and quantitative ΔF/F (bottom) of phasic dopamine activity aligned to the CS+ and CS-cues following training. (F) Color plot of dopamine fluctuations during 15 sec window surrounding presentation of the CS+ and following ICV administration of 0.3 μg or 1 μg oxytocin. Red boxes highlight first ten trials, where oxytocin had greatest effect on phasic dopamine signal. (G) Representative traces (left) and quantitative ΔF/F (right) of cue-evoked phasic dopamine during 15 sec window surrounding presentation of the CS+ and following ICV administration of 0.3 μg or 1 μg oxytocin. (H) Quantitative ΔF/F of first lick-evoked phasic dopamine during 15 sec window surrounding first lick after sipper presentation and following ICV administration of 0.3 μg or 1 μg oxytocin.

## Discussion and Conclusions

Central oxytocin signaling potently reduces food intake and plays a key role in regulating overall energy balance (Blevins and Ho, 2013; Leng et al., 2008). The behavioral and neuronal mechanisms mediating these effects are poorly understood. In order to probe the behavioral intricacies through which oxytocin inhibits food intake, we looked at the effect of central (lateral ICV) oxytocin administration in a number of distinct conditioned palatable food reinforcement-associated behavioral tasks. Results reveal that central oxytocin reduced palatable food seeking behavior in the absence of consumption in a nonreinforced conditioned place preference test. ICV oxytocin also reduced effort-based palatable food-directed operant responding under ad libitum-fed (nonrestricted) conditions in the DRL task, resulting in a higher task efficiency which is indicative of reduced impulsivity. These findings may also result from decreased motivation for sucrose, and thus we also examined the effect of oxytocin in an effort-based decision-making task. We found that ICV oxytocin reduced effort-based operant responding for palatable food under food-restricted conditions in the PROG vs. chow choice task, where oxytocin shifted the ratio of food consumed from effort-based responding (lever pressing) on a PROG schedule for a preferred food (sucrose) to the less preferred lab chow that is freely available in the apparatus. Consistent with a preference shift in favor of chow over sucrose, ICV oxytocin also significantly reduced consumption of sucrose but not chow in a choice test in which both foods were freely available in the training apparatus. Additional results show that central oxytocin signaling did not influence incentive learning, or the process in which rats learn about the value of rewards based on postingestive feedback. Furthermore, oxytocin did not affect general activity levels in an open field test, suggesting that the reductions in sucrose-motivated behavior were not secondary to lethargy or impaired motor responses. Overall these findings show that central oxytocin signaling reduces palatable food-directed responses under various conditions but does not significantly influence learning about the postingestive reinforcing properties of sucrose consumption.

Modulation of phasic dopamine neural responses is a feasible mechanism through which oxytocin reduces palatable food-directed responses, as the conditioned place preference test, DRL impulsivity test, and PROG vs. chow choice test are all influenced by pharmacological manipulations targeting dopamine receptors (Randall et al., 2012; Simon et al., 2013; Spyraki et al., 1982). Moreover, oxytocin neurons in the paraventricular hypothalamic nucleus (PVH) directly project to VTA dopamine neurons (Beier et al., 2015; Peris et al., 2017; Xiao et al., 2017) and oxytocin administration to the VTA suppresses sucrose intake (Mullis et al., 2013). Here we recorded phasic dopamine neuron calcium activity in awake and behaving male rats in an appetitive Pavlovian conditioning task and results revealed that central oxytocin administration suppressed food (sucrose) cue-evoked dopamine neuron responses. These effects were observed during the first ten trials only (within ∼20 min into the session), which suggests that oxytocin-mediated inhibition of food reward seeking behavior, consistent with effects on reducing chow intake (Ho et al., 2014; Liu et al., 2020; Ong et al., 2015), is immediate and short-lasting. By providing an immediate and brief anorexigenic signal, oxytocin may be involved in attention-redirection for need-based prioritization of behavior (e.g., contextual and energetic state-driven shifting between appetitive and social behavior) (Burnett et al., 2019), which is an area for future investigation.

Electrophysiological studies demonstrate that oxytocin administration, as well as optogenetic stimulation of oxytocinergic terminals, enhances activity of VTA dopamine neurons (Xiao et al., 2017). In contrast, systemic oxytocin administration reduces dopamine release (as measured by in vivo fixed potential amperometry) in the nucleus accumbens under baseline conditions and following nomifensine, a dopamine reuptake inhibitor (Estes et al., 2019). Consistent with this latter report, present results reveal reduced dopamine neuron activity in response to presentation of a sucrose-conditioned cue. It is possible these conflicting results occur because, while oxytocin reduces motivated behavior for palatable food, oxytocin enhances social reward and modulates attention-orienting responses to external contextual social cues (Shamay-Tsoory and Abu-Akel, 2016). Indeed, VTA oxytocin administration increases dopamine release in the nucleus accumbens in response to sociality and activation of this circuit enhances prosocial behaviors (Hung et al., 2017; Shahrokh et al., 2010), an effect that occurs through coordinated activity with serotonin (Dolen et al., 2013). Thus, the influence of oxytocin signaling on dopamine neural responses may be dependent on the conditions of behavioral testing, with oxytocin enhancing dopamine signaling in the context of sociality, and reducing dopamine signaling in the context of food-associated stimuli.

In addition to a complex reinforcement-dependent relationship between oxytocin and dopamine signaling, oxytocin has also been proposed to be conditionally anorexigenic depending on social context (Olszewski et al., 2016). For example, intranasal oxytocin increases food sharing in the common vampire bat (Carter and Wilkinson, 2015) and peripheral oxytocin enhances the social transmission of food preference in rats (Popik and Van Ree, 1993). Moreover, hierarchy social factors influence the effectiveness of oxytocin receptor ligands on food intake. Specifically, administration of an oxytocin receptor antagonist in dominant mice increases the amount of sugar consumed regardless of social or nonsocial contexts, whereas in subordinate rats, it only increases the amount of sugar consumed in a setting devoid of social cues (Olszewski et al., 2015).

In contrast to the enhancement of social reinforcement and similar to its effects on reducing food intake and food-motivated responses, oxytocin also potently inhibits ethanol, methamphetamine, and cocaine seeking and consumption (Cox et al., 2017; Kohtz et al., 2018; Peters et al., 2017). Oxytocin has been proposed to promote social affiliation, in part, by reducing behavioral effects of drugs of abuse via alterations in dopamine neurotransmission (Insel, 2003; Mattson et al., 2001), a model that can potentially apply to reduction of food-motivated behaviors. Consistent with this framework, suckling stimulation in lactating dams and cocaine exposure in virgin females activate the dopamine system. However, lactating dams exposed to cocaine show reduced activity in dopaminergic-associated pathways, which the authors suggest may be in part due to oxytocin (Ferris et al., 2005). These studies collectively suggest that oxytocin signaling may prioritize behavior towards socially-relevant cues in favor or other reinforcers. Future studies are needed to further examine whether oxytocin’s influence on dopamine signaling and food-motivated behavior are dependent on the nature of the reinforcement and the context.

Our findings demonstrate that while dopamine-associated conditioned food reward-directed behaviors were reduced under various conditions following ICV oxytocin administration, central oxytocin did not influence incentive learning. Specifically, ICV oxytocin administered prior to consuming sucrose after an energetic motivational shift (from satiety to hunger/fasted) did not affect sucrose-motivated behavior when subsequently tested after a fast. When considered together with our results showing that central oxytocin reduced dopamine neuron responses to a sucrose-associated cue, these results are consistent with previous literature demonstrating that instrumental incentive learning, using procedures similar to those in the present study, is not affected by treatment of flupenthixol, a D1 and D2 receptor antagonist (Wassum et al., 2011b). Furthermore, our incentive learning results suggest that oxytocin may act downstream via a ventral and not dorsal striatal pathway to influence food-motivated behavior. For example, Tellez and colleagues identified dissociable basal ganglia sensorimotor circuits that encode hedonic versus metabolic values of reward (Tellez et al., 2016), where dorsal striatal descending pathways are recruited to encode the nutritional (postingestive) aspects of food, and ventral striatum descending pathways transmit hedonic (flavor) value of food. These findings combined with present results support a putative model in which oxytocin signals downstream to ventral striatal dopaminergic pathways to reduce conditioned food cue-directed motivated behaviors, without influencing dorsal striatum-mediated postingestive incentive learning.

While present results identify a role for central oxytocin signaling in modulating palatable food cue-directed behavior and mesolimbic dopamine signaling, one limitation of the present study is that the mediating sites of CNS oxytocin action were not investigated. Oxytocin receptors are expressed widely in various feeding-associated regions throughout the brain, including in the amygdaloid complex, VTA, hippocampus, basal ganglia, prefrontal cortex, caudal brainstem, and hypothalamus (Freund-Mercier et al., 1987; Otero-Garcia et al., 2016). In addition to the VTA, one region that may be relevant to present results is the ventral subregion of the hippocampus, which has robust oxytocin receptor expression (Freund-Mercier et al., 1987) and is involved in food impulsivity (Noble et al., 2019), conditioned place preference for palatable food (Kanoski et al., 2011a; Zhou et al., 2019), and striatal dopamine signaling (Kanoski et al., 2013). Additionally, the prefrontal cortex plays a critical role in effort-related food choice behavior (Walton et al., 2002) as well as food impulsivity (Hsu et al., 2018b), and may therefore be a likely candidate site of action for oxytocin’s effects on food reward-directed behavior. In addition to these forebrain regions, oxytocin signaling potently reduces feeding in the hindbrain nucleus tractus solitarius (mNTS). While this region is more classically associated with meal size and satiation control, recent results link the mNTS in the control of conditioned food reward-directed behaviors (Kanoski et al., 2014a; Liu and Kanoski, 2018). These and other potential sites of action should be explored in future studies, as they may act in conjunction and/or in parallel with dopaminergic mesolimbic ventral striatal pathways to mediate oxytocin’s suppression of palatable food cue-directed responses. A second limitation of the present study is that, with the exception of the photometry experiment, we utilized a single dose (1 μg) of oxytocin for behavioral analyses. While this dose was selected based on our previous work where it reliably reduced 30 min chow intake in rats following ICV administration (Liu et al., 2020), oxytocin has been shown to have dose-related U-shaped relationship with social trust behavior in humans (Zhong et al., 2012). Thus, a future direction is to examine whether such U-shaped relationships occur when various doses of central oxytocin are tested on conditioned food reward-directed behaviors. Lastly, with regards to the photometry, the present study recorded dopamine neuron activity in the VTA, however, it is unclear if this activity resulted in dopamine release and, if so, in which terminal regions (e.g. nucleus accumbens, medial prefrontal cortex, etc.). A number of recent investigations have demonstrated dopamine release can occur independent of cell body activity (Mateo et al., 2017; Mohebi et al., 2019; Threlfell et al., 2012). Future work will need to utilize fluorescent dopamine sensors (dlight, GRAB-DA) to determine downstream dopamine release dynamics in response to central oxytocin signaling.

Collectively, our results reveal that central oxytocin administration reduces motivated behaviors in response to conditioned food cues under various conditions. Furthermore, oxytocin suppresses food cue-evoked phasic VTA dopamine neuron responses, thus providing a potential neurobiological mechanism for the capacity of central oxytocin signaling to rapidly inhibit palatable food-directed behavior. These results illuminate novel behavioral and neurochemical mechanisms through which oxytocin inhibits food intake, thus further supporting oxytocin’s potential as a target system for obesity pharmacotherapy development.

## Acknowledgements

We thank Elizabeth Davis, Sarah Terrill, Linda Tsan, and Lekha Chirala for their critical contributions to the research. This study was supported by the National Institute of Health grants: DK118402 (SEK), DK104897 (SEK), DA025634 (MFR), DK111158 (EEN), DA047052 (TMH), DK118944 (CML), DK116558 (ANS).

